## Supplementary data file for "Human serum triggers antibiotic tolerance in *Staphylococcus* aureus"

**Figures S1-S9**

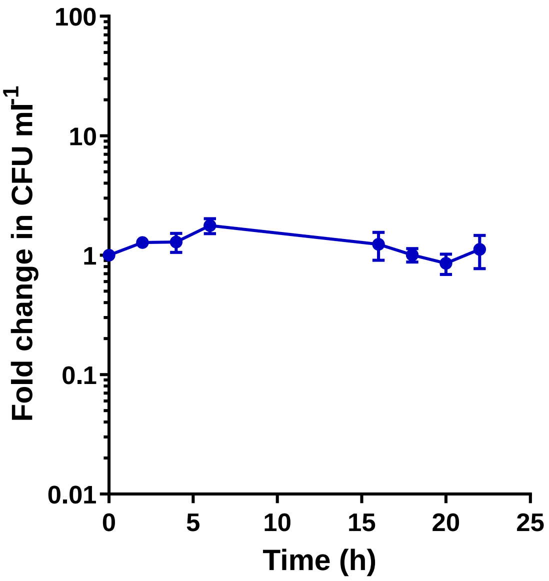


**Figure S1. *S. aureus* CFU counts do not change during incubation in human serum.** Fold change in CFU ml$^{-1}$ during incubation of TSB-grown *S. aureus* in human serum for 22 h. Graph represents the mean ± standard deviation of three independent experiments. Data were analysed by one-way ANOVA and no statistically significant differences were observed.

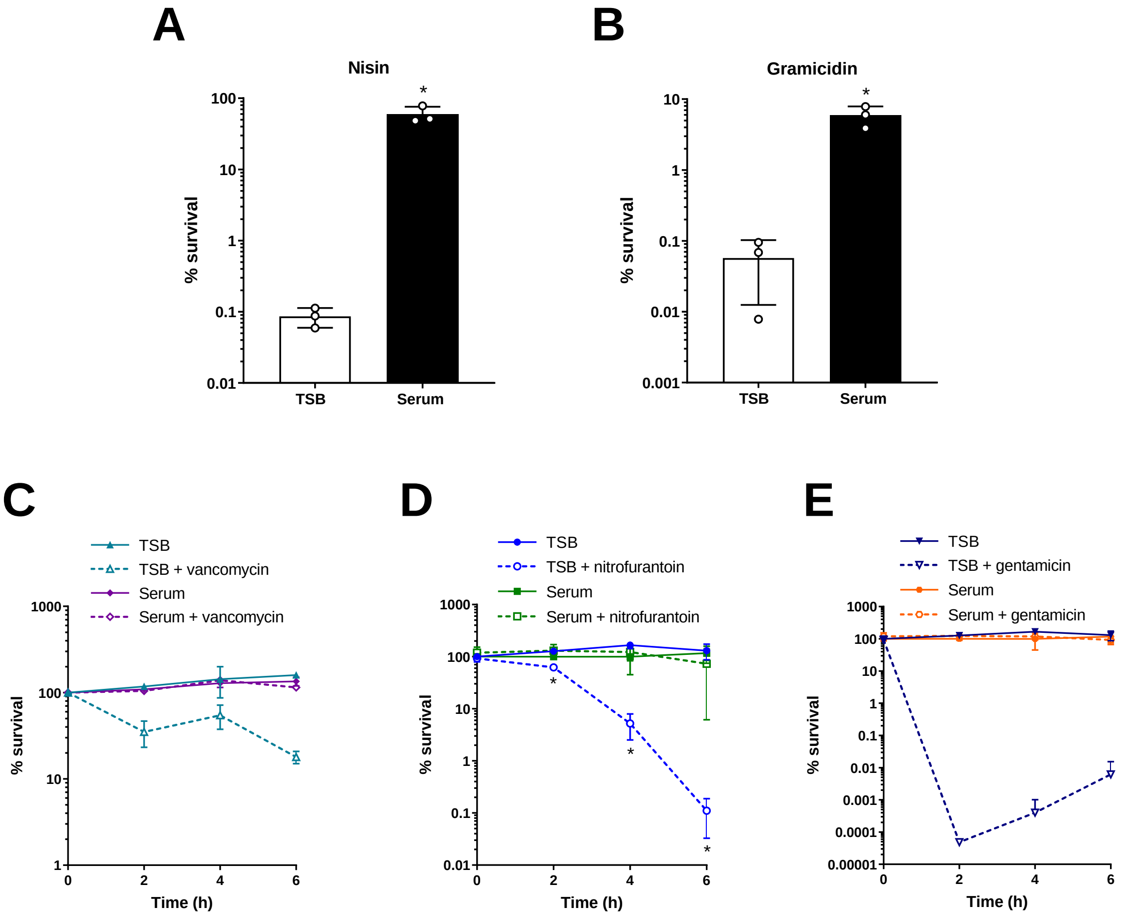


**Figure S2. Incubation of *S. aureus* in serum results in tolerance towards antimicrobials from various classes.** Percentage survival of TSB-grown and serum-adapted cultures of *S. aureus* USA300 WT after/throughout a 6 h incubation in serum with 128 µg ml$^{-1}$ nisin (**A**), 160 µg ml$^{-1}$ gramicidin (**B**), 160 µg ml$^{-1}$ vancomycin (**C**), 160 µg ml$^{-1}$ nitrofurantoin (**D**) or 40 µg ml$^{-1}$ gentamicin (**E**). Graphs represent the mean ± standard deviation of three independent experiments. Graphs in **A** – **B** were analysed by t-test (TSB-grown vs serum-adapted). *, $P < 0.05$. Graphs in **C** – **E** were analysed by two-way ANOVA with Tukey's *post-hoc* test, TSB-grown compared to serum-adapted at each time-point).

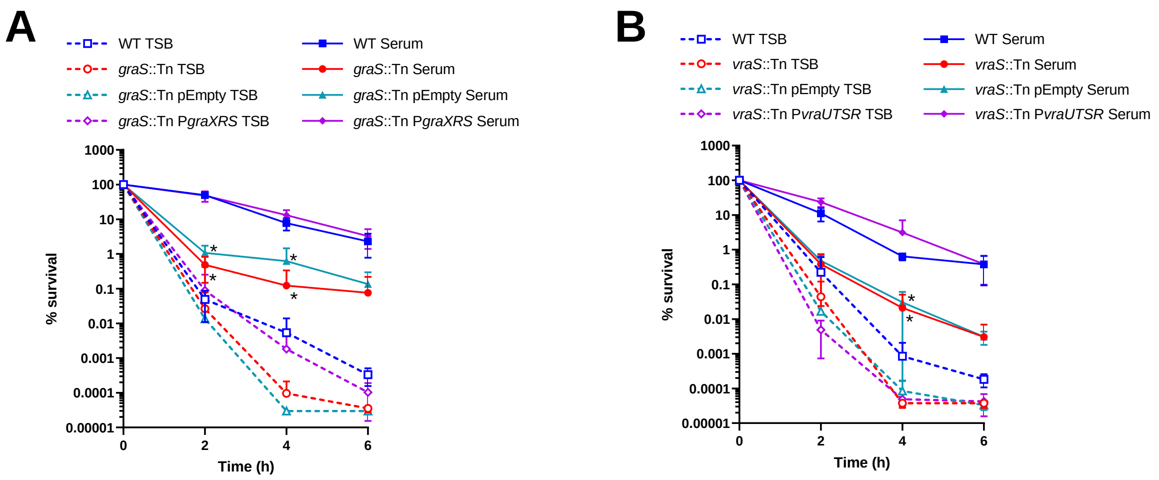


**Figure S3. Complementation of the *vraS*::Tn and *graS*::Tn mutants restores tolerance to WT levels.** Percentage survival of TSB-grown and serum-adapted cultures of *S. aureus* JE2 WT, *graS*::Tn and *graS*::Tn complemented with pCN34 or P*graXRS* **(A)** and WT, *vraS*::Tn and *vraS*::Tn complemented with empty pCN34 or P*vraUTSR* **(B)** over 6 h incubation in serum with 80 µg $ml^{-1}$ daptomycin. Graphs represent the mean ± standard deviation of three independent experiments. *, $P < 0.05$ (two-way ANOVA with Dunnett's *post-hoc* test, serum-adapted WT compared to serum-adapted mutants at each time-point).

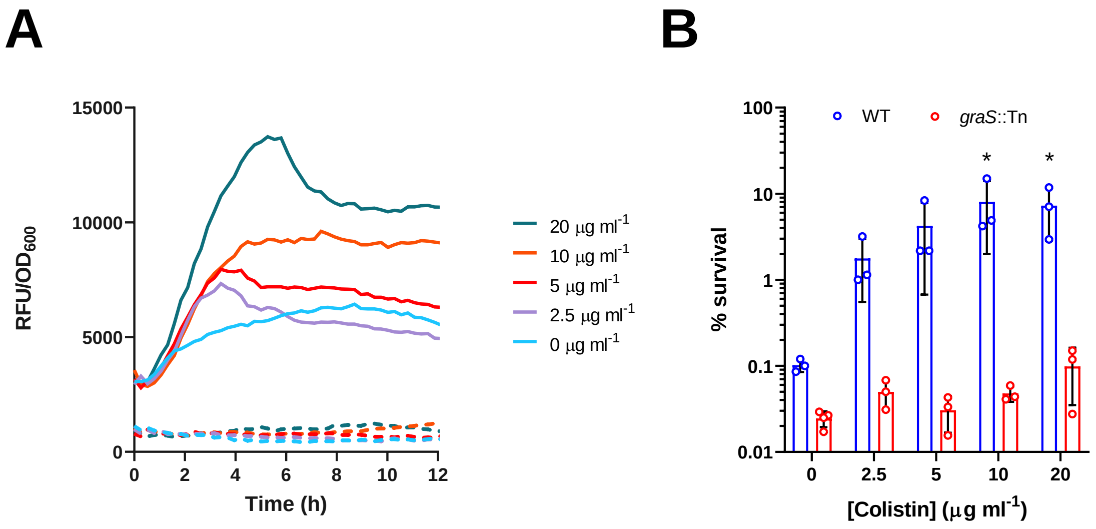


**Figure S4. Colistin activates GraRS signalling and triggers daptomycin tolerance.** TSB-grown cultures of *S. aureus* JE2 WT (solid lines) and the *graS*::Tn mutant (dashed lines) containing P*dltA*-*gfp* were exposed to various concentrations of colistin (2.5 – 20 µg $ml^{-1}$) in RPMI 1640 (**A**). GFP fluorescence (RFU) and $OD_{600}$ were measured every 15 min for 12 h. Fluorescence values were divided by $OD_{600}$ measurements to normalise for changes in cell density. Percentage survival after 6 h exposure to 80 µg $ml^{-1}$ daptomycin of *S. aureus* JE2 WT and the *graS*::Tn mutant which had been pre-incubated for 16 h in RPMI 1640 only or RPMI 1640 supplemented with indicated concentrations of colistin (**B**). Graph in **A** represents the mean of three independent experiments and error bars have been omitted for clarity. Graph in **B** represents the mean ± standard deviation of three independent experiments. *, $P < 0.05$ (two-way ANOVA with Dunnett's *post-hoc* test, RPMI 1640 and colistin compared to RPMI 1640 only). RFU, relative fluorescent units; $OD_{600}$, optical density at 600 nm.

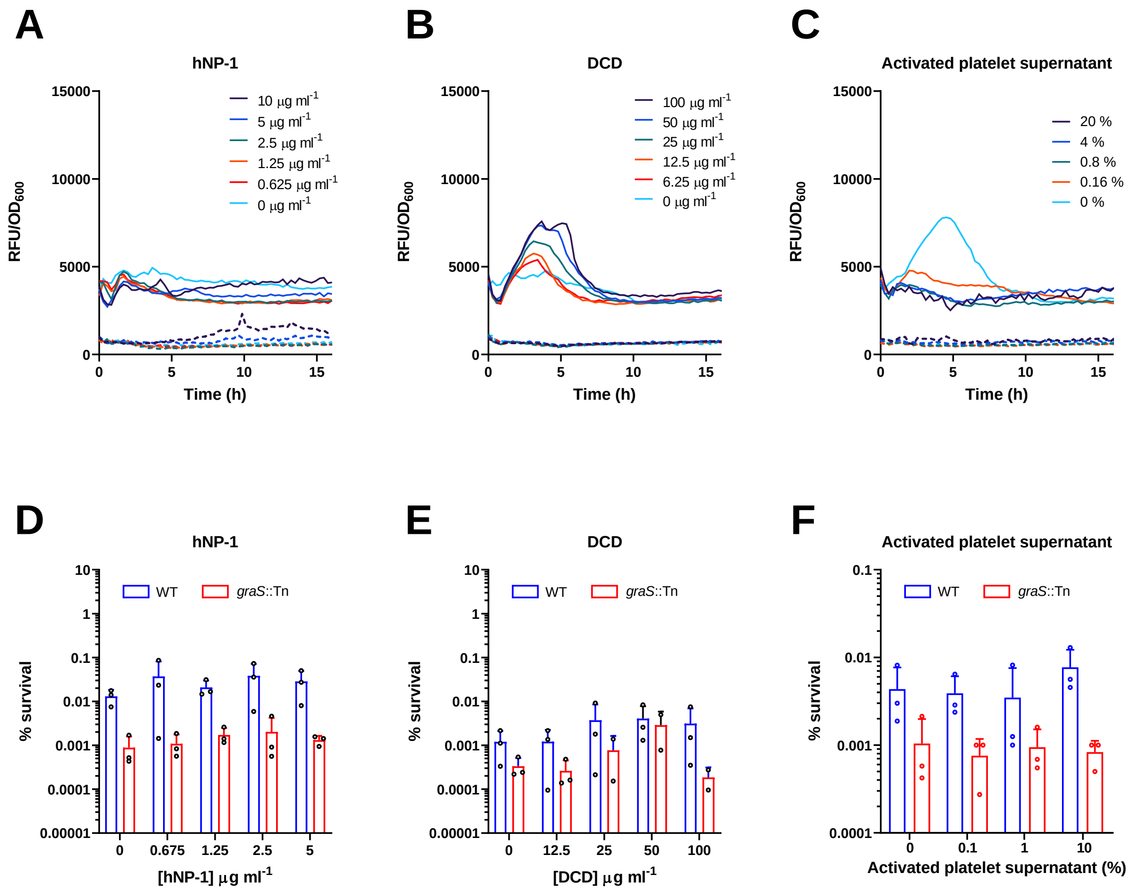


**Figure S5. Human neutrophil peptide-1, dermcidin and platelet AMPs do not activate GraRS signalling or induce daptomycin tolerance.** TSB-grown cultures of *S. aureus* JE2 WT (solid lines) and the *graS*::Tn mutant (dashed lines) containing P*dltA-gfp* were exposed to a range of concentrations of hNP-1 (**A**), DCD (**B**) or activated platelet supernatant (**C**) in RPMI 1640 and GFP fluorescence (RFU) and $OD_{600}$ were measured every 15 min for 16 h. Fluorescence values were divided by $OD_{600}$ measurements to normalise for changes in cell density. Percentage survival after 6 h exposure to 80 µg $ml^{-1}$ daptomycin of WT and *graS*::Tn mutant strains which had been incubated for 16 h in RPMI supplemented with hNP-1 (**D**), DCD (**E**) or activated platelet supernatant (**F**). Graphs in **A** – **C** represent the mean of three independent experiments with error bars omitted for clarity. Graphs in **D** – **F** represent the mean ± standard deviation of three independent experiments. Data in **D** – **F** were analysed by two-way ANOVA. No statistically significant differences were observed (RPMI 1640 + AMPs compared to RPMI 1640 alone).

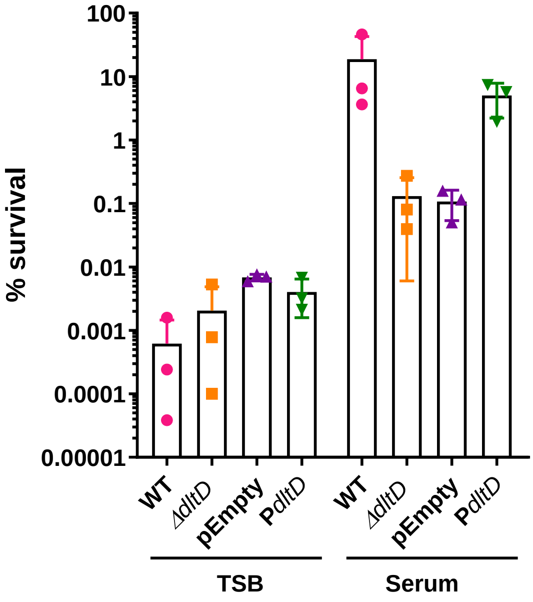


**Figure S6. Complementation of *ΔdltD* mutant with WT *dltD* restores tolerance to WT levels.** Percentage survival of TSB-grown and serum-adapted cultures of WT, the *ΔdltD* mutant or *ΔdltD* mutant complemented with either pCN34 or P*dltD* after a 6 h daptomycin exposure in serum. Graph represents the mean ± standard deviation of three independent experiments. No statistically significant differences were observed (one-way ANOVA, serum-adapted WT compared to serum-adapted mutants).

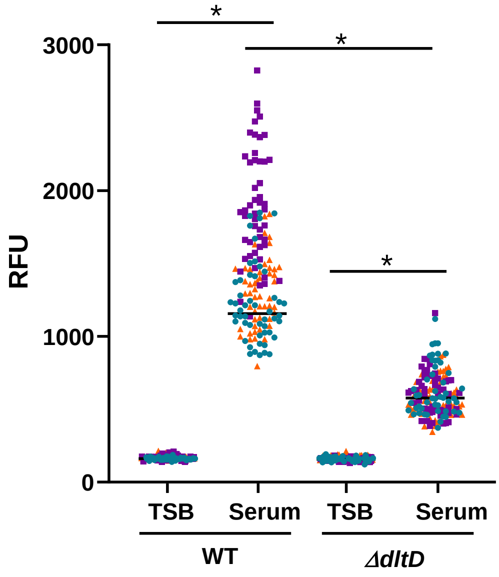


**Figure S7. The *ΔdltD* mutant incorporates less HADA than WT during serum adaptation.** The fluorescence of individual TSB-grown and serum-adapted WT and *ΔdltD* cells was quantified. Graph represents the fluorescence of 50 cells per biological replicate (150 cells in total) with the mean of the three replicates indicated. Each biological replicate is depicted in a different colour. Data were analysed by Kruskal Wallis test (*, $P < 0.05$).

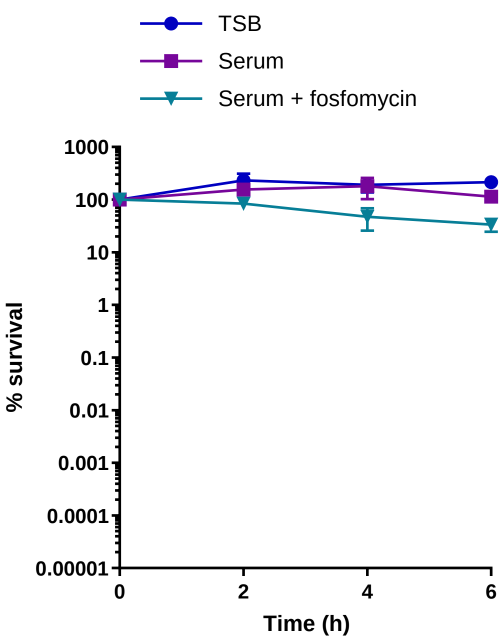


**Figure S8. Inhibition of peptidoglycan synthesis during serum adaptation does not affect bacterial viability.** Percentage survival over 6 h of TSB-grown *S. aureus* or cultures which had been incubated in serum supplemented, or not, with 64 µg ml$^{-1}$ fosfomycin for 16 h. Graph represents the mean ± standard deviation of three independent experiments. No statistically significant differences were observed (two-way ANOVA, serum-adapted vs serum + fosfomycin-adapted).

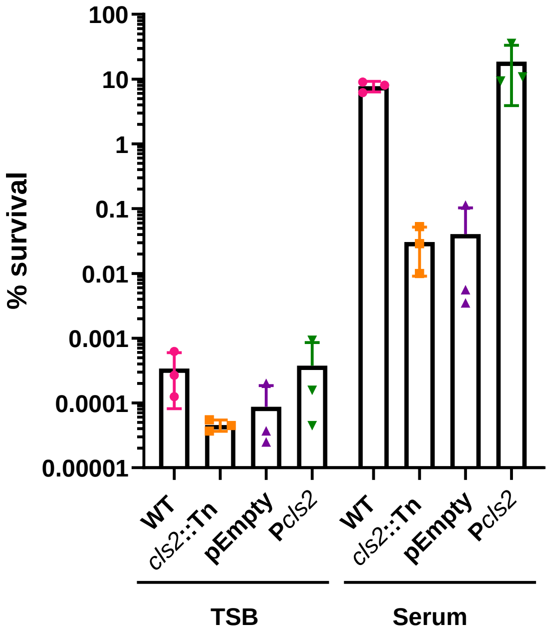


**Figure S9. Complementation of the *cls2*::Tn mutant restores daptomycin tolerance.** Percentage survival of TSB-grown and serum-adapted cultures of *S. aureus* JE2 WT, *cls2*::Tn or *cls2*::Tn complemented with empty pCN34 or P*cls2* after a 6 h exposure to 80 µg ml$^{-1}$ daptomycin in serum. Graph represents the mean ± standard deviation of three independent experiments. No statistically significant differences were observed (one-way ANOVA, serum-adapted WT compared to serum-adapted mutants).
